# Intraspecies sequence-graph analysis of the *Phytophthora theobromicola* genome reveals a dynamic structure and variable effector repertoires

**DOI:** 10.1101/2025.07.17.665439

**Authors:** Jadran F García, Rosa Figueroa-Balderas, Michael E.H. Matson, Shahin S. Ali, Bryan A. Bailey, Jean-Philippe Marelli, Dario Cantu

## Abstract

*Phytophthora theobromicola* is an emerging cacao pathogen recently identified in Brazil as an aggressive agent of black pod rot. We generated genome assemblies for two *P. theobromicola* isolates using long-read sequencing and five additional isolates using short reads. Comparative analysis revealed a genome size and predicted gene content comparable to *P. citrophthora*, a closely related species with a broad host range that includes both citrus and cacao. An intraspecies sequence-graph analysis revealed a highly dynamic genome structure with high proportion of variable effectors. Syntenic orthology analysis across 13 *Phytophthora* species identified orthologous gene groups conserved only in cacao pathogens and others specific to *P. theobromicola*. RxLR effectors and CAZymes were particularly enriched among lineage-specific syntenic groups, with RxLRs preferentially located near transposable elements and within gene-sparse, repeat-rich regions. Transcriptome analysis of infected cacao tissues showed that 88% of predicted effectors were expressed, with pods exhibiting the highest number of upregulated genes. Notably, several RxLRs classified as *P. theobromicola-*specific syntenic orthologs were highly expressed in infected tissues, suggesting that these lineage-specific effectors may play key roles in host-pathogen interactions unique to cacao. Together, our findings highlight the dynamic architecture and functional plasticity of the *P. theobromicola* genome, providing foundational insights into its virulence strategies and supporting future studies on host adaptation and effector evolution in emerging cacao pathogens.

## Introduction

The worldwide chocolate industry, valued at over 130 billion US Dollars annually (Zion Market Research faces significant treats from multiple cacao diseases that reduce both yield and quality (Marelli et al. 2019). It is estimated that up to 38% of global annual cacao bean production is lost to disease, with where more than half of these losses attributed to *Phytophthora* species (Marelli et al. 2019). Members of this genus can infect all parts of the cocoa plant, causing black pod rot, stem canker, and leaf blight (Firman and Vernon 1970; Marelli et al. 2019; Appiah et al. 2004). Among the major cacao diseases, black pod disease (also known as black pod rot) is particularly devastating, causing yield losses of 20–30% on average, with some farmers in wetter regions reporting complete crop failure (Drenth and Guest 2013). The lesions caused by the disease can appear at any stage of development and at any part of the pod, starting as small, hard, dark spots and growing rapidly to cover the entire pod damaging the beans inside (Guest 2007). The progression of the disease is influenced by multiple factors, including the *Phytophthora* species, cacao genotype, humidity, rainfall, and temperature (Guest 2007; Puig et al. 2018; de Oliveira and Luz 2005).

The development of disease is facilitated by an arsenal of proteins secreted by the pathogen that interact with host cells (McGowan and Fitzpatrick 2017; Kamoun 2006). These proteins, collectively referred to as effectors, are broadly classified by their site of action: apoplastic effectors act in the extracellular space, while cytoplasmic effectors are translocated into the host cell. Apoplastic effectors include toxins such as necrosis- and ethylene-inducing peptide 1-like proteins (NLPs), which have been shown to trigger cell death in model plants. In *Phytophthora* species, NLPs are associated with disruption of host membrane integrity (Lenarčič et al. 2017; Yan Wang and Wang 2018). The apoplastic compartment also contains a diverse set of hydrolytic enzymes that degrade plant cell wall components, facilitating nutrient acquisition. Among these, carbohydrate-active enzymes (CAZymes)—including cutinases, pectinases, and glycoside hydrolases are enriched in *Phytophthora* genomes (Jiang and Tyler 2012; Yan Wang and Wang 2018; Ma et al. 2015). Among cytoplasmic effectors, two major families have been well characterized in *Phytophthora*: RxLR and Crinkler (CRN) effectors (McGowan and Fitzpatrick 2017; Whisson et al. 2007; Schornack et al. 2010). RxLR effectors are defined by an N-terminal motif containing “RxLR” and “EER” sequences, which are necessary for host cell translocation (McGowan and Fitzpatrick 2017; Jiang and Tyler 2012; Yan Wang and Wang 2018). In some cases, this translocation can occur independently of additional pathogen-encoded machinery (Dou et al. 2008; H. Wang et al. 2023). Functionally, RxLR effectors have been shown to either suppress host programmed cell death or, conversely, induce it, depending on the host context and specific effector (Q. Wang et al. 2011). Crinklers (CRNs) are characterized by an “LxLFLAK” motif that facilitates translocation (McGowan and Fitzpatrick 2017; Schornack et al. 2010) and are known to localize to the host nucleus (Schornack et al. 2010; Stam et al. 2013). CRNs have been implicated in both induction and suppression of host cell death (Liu et al. 2011; Stam et al. 2013), although the specific mechanisms are not yet fully understood. Both RxLRs and CRNs are often located in gene-sparse, repeat-rich regions of the genome, which are associated with accelerated evolution and diversification of virulence factors (Stassen and Van den Ackerveken 2011).

At least five *Phytophthora* species have been reported to cause black pod rot with significant commercial impact: *P. palmivora, P. megakarya*, *P. capsici*, *P. citrophthora*, and, more recently, *P. theobromicola* (Marelli et al. 2019; Decloquement et al. 2021; Surujdeo-Maharaj et al. 2016). Both *P. palmivora* and *P. megakarya* have undergone recent and independent whole-genome duplications, with *P. megakarya* possessing one of the largest known *Phytophthora* genomes at approximately 222 Mbp (Morales-Cruz et al. 2020). This duplication contributed to a substantial increase in gene content, with an average of 57,577 protein-coding genes in *P. megakarya* and 36,778 in *P. palmivora*. These expansions include a high number of putative virulence factors, particularly effector proteins such as RxLRs, CRNs, and NPP1-like proteins (a class of NLP), underscoring the role of gene duplication and structural variation in effector diversification and pathogen adaptation.

*Phytophthora theobromicola* sp. nov was recently isolated from a set of isolates obtained from cacao pods exhibiting symptoms of black pod disease (Decloquement et al. 2021). The identification of the new species was based on a combination of morphological features and multilocus phylogenetic analysis. Molecular markers used for this analysis included sequences from the β-tubulin, elongation factor 1-alpha, heat shock protein 90, internal transcribed spacer (ITS), and cytochrome c oxidase subunits I and II genes, which together supported the designation of *P. theobromicola* as a distinct species. *Phytophthora theobromicola* isolates were found to be more aggressive than *P. palmivora*, suggesting a high virulence potential (Decloquement et al. 2021).

In response to its recent emergence and demonstrated pathogenicity, we present here the first genomic resources for *P. theobromicola*, providing a foundation for future studies on its evolutionary history, adaptive potential, and virulence mechanisms. We assembled high-quality diploid genomes for two isolates using long-read sequencing and generated short-read assemblies for five additional isolates. We performed functional annotation of candidate effectors and constructed a pangenome graph including all isolates to assess intraspecific variability in effector content. Comparative genomic analysis with multiple *Phytophthora* species revealed groups of syntenic orthologs shared among cacao-infecting species, as well as orthologs unique to *P. theobromicola*. To characterize gene expression dynamics during infection, we analyzed RNA-seq data from inoculated cacao tissues, revealing the large set of predicted effectors expressed during pod infection.

## Methods

### Inoculation and nucleic acid extraction

*Phytophthora theobromicola* isolates obtained from symptomatic pods of different cultivation sites in Bahia, Brazil were used for this study (Decloquement et al. 2021). Mycelial plugs from a 5-day-old culture grown on carrot-agar (CA) medium were used to inoculate cacao pods, leaves, and stems. Control inoculations were made using CA medium plugs with no mycelial growth as described in Decloquement et al., (2021). The genomic DNA (gDNA) for short-read sequencing, high molecular weight (HMW) gDNA for SMRT sequencing, and the total RNA from the inoculated tissues and the pure mycelia was extracted as previously described in Morales-Cruz et al. (2020).

### Sequencing library preparation

The high molecular weight (HMW) gDNA of *Phytophthora theobromicola* P0449 and MB01960 was fragmented using a 26G blunt needle (SAI Infusion Technologies, IL, USA) by aspirating the entire volume and passing the sample through the needle fifteen times. After shearing, the sample was cleaned and concentrated using 0.45× AMPure PB beads. The size distribution of the sheared gDNA fragments was evaluated using pulsed-field gel electrophoresis (Pippin Pulse, Sage Science, MA, USA) prior to library preparation. Continuous Long Read (CLR) libraries were prepared using the SMRTbell Express Template Prep Kit 2.0 (Pacific Biosciences, CA, USA) following the manufacturer’s instructions. Up to 5 µg of SMRTbell template was size-selected with the Sage Blue Pippin (Sage Science, MA, USA) using a cutoff range of 17–80 kb. The size-selected library was cleaned with 1× AMPure PB beads and sequenced on the PacBio Sequel II platform (DNA Technology Core Facility, University of California, Davis). Likewise, The genomic DNA of the remaining *P. theobromicola* isolates was sheared to approximately 450 bp using a Covaris E220 sonicator (PerkinElmer, MA, USA). DNA-seq libraries were prepared from the sheared DNA using the KAPA LTP Library Preparation Kit (Roche Diagnostics, IN, USA). The libraries were sequenced in 150 bp paired-end mode (2 x150) on an Illumina HiSeq 4000 (Novogene Co. Ltd., Beijing, China).

The RNA extracted from the pure mycelium, inoculated tissue and control tissue was used to prepare the RNA-seq libraries. A total of 500 ng of RNA per sample was used as input prepare the libraries using the TruSeq RNA Sample Preparation Kit v2 (Illumina, CA, USA), following the manufacturers’ protocols with individual barcoding. Final libraries were assessed for quantity and quality using a High Sensitivity chip on a Bioanalyzer 2100 (Agilent Technologies, CA, USA) and a Qubit fluorometer (Life Technologies, CA, USA). The RNA-seq libraries were sequenced in 150 bp paired-end mode (2 x150) by IDSeq on an Illumina NovaSeq 6000.

### Genome assembly

The assembly of *P. theobromicola* isolates MB01960 and P0449 were assembled using SMRT reads with FALCON-Unzip ver. 2017.06.28-18.01 (Chin et al. 2016) using a custom pipeline published in (Minio et al. 2019) and available at https://github.com/andreaminio/FalconUnzip-DClab. Different parameter combinations were tested to obtain the least fragmentated assembly as described in (Morales-Cruz et al. 2020). Haplotype phasing was performed with default parameters (Chin et al. 2016). Primary contigs and haplotigs were polished with Arrow (from ConsensusCore2 v.3.0.0) using raw long reads. Primary contigs and haplotigs were scaffolded using SSPACE-Longreads v.1.1 (Boetzer and Pirovano 2014). The genomes of the isolates in **Supplementary table 1** were assembled using paired-end short read sequencing data. Raw reads were quality-filtered and adapter clipped using Trimmomatic v.0.36 (Bolger, Lohse, and Usadel 2014), with the following settings: ILLUMINACLIP:2:30:10 LEADING:7 TRAILING:7 SLIDINGWINDOW:10:20 MINLEN:36. SPAdes v4.0.0 (Prjibelski et al. 2020) was used to assemble the quality-filtered reads with the careful option and automatic read coverage cutoff after optimizing the multiple kmer combination (-k 55,77,99,111,127). These contigs were labeled “main” if their length was at least 1,000 bp and “short” for contigs smaller than 1,000 bp. To assess the assembly completeness of the genomes, we performed Benchmarking Universal Single-Copy Orthologs (BUSCO v.5.4.2;(Manni et al. 2021) analysis with the stramenopiles_odb10 lineage dataset.

### Repeat and gene annotation

The concatenated genomes of all *Phytophthora* species in **Table 1** were used to predict repeat models using RepeatModeler v. 2.0.5 (Flynn et al. 2020; Smit, Hubley, and Green 2015) with default parameters and -LTRStruct option to predict LTR retrotransposons based on their structure. The predicted models were concatenated with the RepeatMasker-RepBase database (release 20181026) and used with RepeatMasker v.4.1.5 (Smit, Hubley, and Green, n.d.) to mask the repeats in each *P. theobromicola* genome. The tool maskFastaFromBed (Quinlan 2014) was used to softmask the repeats in all the genomes.

**Table 1.** Genome assembly and gene annotation statistic of *P. theobromicola* primary assembly and other *Phytophthora* species affecting *Theobroma cacao*.

| Species | *P. theobromicola*<br>(MB01960) | *P. theobromicola*<br>(P0449/ATCC) | *P. megakarya*<br>(PmGH34) | *P. palmivora*<br>(PpGH49) | *P. capsici*<br>(LT1534) | *P. citrophthora*<br>(STE-U-9442) |
| --- | --- | --- | --- | --- | --- | --- |
| Haploid assembly size (Mb) | 47.5 | 45.6 | 214.2 | 115.5 | 94.2 | 48.5 |
| Coverage | 757 x | 767 x | > 150 x | > 150 x | 35 x | 521 x |
| No. of scaffolds | 52 | 27 | 502 | 240 | 782 | 155 |
| N50 (Mb) | 1.9 | 2.1 | 0.8 | 0.9 | 0.5 | 0.9 |
| L50 (scaffold no.) | 11 | 7 | 78 | 43 | 44 | 16 |
| BUSCOs in assembly (%) | 100 | 96 | 100 | 100 | 100 | 100 |
| Repeats masked (Mb) | 13.7 | 12.8 | 139 | 56.6 | 46.4 | 14.4 |
| Repeats masked (%) | 28.8 | 28.1 | 64.9 | 49 | 49.3 | 29.6 |
| No. of CDS | 16,387 | 15,771 | 53,476 | 31,821 | 23,748 | 16,447 |
| Mean protein size | 471 | 469 | 340 | 427 | 434 | 498 |
| BUSCOs in gene prediction (%) | 98 | 96 | 95 | 95 | 99 | 91 |
| Mean gene density (gene/10 kb) | 4.0 | 4.0 | 2.8 | 3.1 | 2.8 | 3.9 |
| Source | This work | This work | Morales-Cruz et al., 2020 | Morales-Cruz et al., 2020 | (Stajich et al. 2021) | (Möller et al. 2024) |

The RNAseq data obtained from pure mycelium and inoculated material was quality filtered and mapped to all softmasked *P. theobromicola* genomes individually using Hisat2 v.2.2.1 (Kim et al. 2019) with --dta --very-sensitive options. The gene prediction *P. theobromicola* genomes was made with Braker2 v2.1.6(Stanke et al. 2006; 2008; Lomsadze, Burns, and Borodovsky 2014; Barnett et al. 2011; Li et al. 2009; Ter-Hovhannisyan et al. 2008; Iwata and Gotoh 2012; Gotoh 2008; Buchfink, Xie, and Huson 2015; Lomsadze et al. 2005; Brůna, Lomsadze, and Borodovsky 2020; Hoff et al. 2019; Brůna et al. 2021; Hoff et al. 2016) using as evidence the mapped RNAseq reads (bam file) and a custom protein database using as base OrthoDB v.11 (Kuznetsov et al. 2023) and predicted proteins of *P. megakarya*, *P. palmivora*, *P. citrophthora* and *P. capsici*. This annotation was cleaned to remove proteins with internal stop codons and without stop codons.

To enhance gene model consistency across all the assemblies and mitigate the effects of fragmentation of some genomes, we performed gene porting by projecting a curated, non-redundant gene set derived from *P. theobromicola* MB1960 on all genomes. The set of non-redundant genes were obtained using the coding sequences of *P. theobromicola* MB1960 with Orthofinder 2.5.5 (Emms and Kelly 2019) and filtering the each orthogroups to obtain the longest representative. This non-redundant set was used to create artificial reference genome and gff3 from the original *P. theobromicola* MB1960. The artificial reference was used with LiftOn v1.0.5 (Shumate and Salzberg 2021; Chao et al. 2025) and each *P. theobromicola* assemblies with the options “-a 0.9 -s 0.9 -exclude_partial -copies -sc 0.95 -polish -cds”. The new lifted annotation was cleaned to remove proteins with internal stop codons (identified with the character *) and proteins without stop codon. The lifted annotation was intersected against the initial braker annotation and the braker genes with no overlap with the lifted genes were kept in the final annotation of each genome. BUSCO v.5.4.2 (Manni et al. 2021) with the stramenopiles_odb10 lineage dataset and the protein mode was used to asses completeness.

### Functional and Effector annotation

The functional annotation was made focusing on effectors and therefore the first task was to identify the secretome. SignalP3 (Dyrløv Bendtsen et al. 2004) was selected for this task over newer versions because previous studies have found that it is the most sensitive at detecting oomycete signal peptides (Sperschneider et al. 2015; McGowan and Fitzpatrick 2017). The mature proteins were used as input for TMHMM v2.0 (Krogh et al. 2001). Proteins were classified as secreted if they had a signal peptide and no transmembrane domain in the first 60 amino acids or more than two transmembrane domains predicted by TMHMM. Then all the secreted proteins were checked with EffectorO (Nur, Wood, and Michelmore 2023) and its machine learning models to predict potential effectors. Next, all the proteins were annotated with the InterProScan v.5.69 pipeline and the databases Pfam, PANTHER, SUPERFAMILY and NCBIfam. To annotate CAZymes, dbCAN3 (Zheng et al. 2023) with the options “--dia_eval 1e-102 --hmm_eval 1e-15 --hmm_cov 0.35” was run locally on all proteins. Annotations validated by two of three tools in dbCAN3 were kept as CAZymes. The R package effectR v.1.0.2 (Tabima and Grünwald 2019) was used to annotate RxLR and Crinkler in all the genomes. First the “regex.search” was used to obtain a training list of RxLR and Crinkler containing the well-known domains, then, a hmm model was trained based on these lists and the proteins were compared against the model to obtain a non-redundant set of putative RxLR and Crinkler effectors. Next, if the protein was classified as secreted and had a functional annotation (in any of the databases with any of the annotation methods used) consistent with known effectors (based on (McGowan and Fitzpatrick 2017)) they were considered putative effectors. **Supplementary table 2** presents the functional annotation with all the methods. **Supplementary table 3** contains the secretome with the putative effector proteins corresponding functional annotation used in the rest of the analyses.

### Sequence graph of effector variability

The genomes of the seven isolates of *P. theobromicola* (including haplotigs) were used to build the sequence graph with the Nextflow v.24.10.2 pipeline nf-core/pangenome (v1.1.2-g0e8a387; https://github.com/nf-core/pangenome/tree/1.1.2; 10.5281/zenodo.10869589; Heumos et al. 2024). The defaults parameters were used except the “wfmash_segment_length” that was set to 3000 based on a previous optimization. The resulting sequences graph was used to classify the nodes in 3 categories. The core nodes were those present in all the genomes, the dispensable nodes were those in more than one genome but not in all of them, and the private nodes were those present exclusively in a single genome. The total set of nodes is considered the full repertoire. The gene classification was based on the node composition. If the majority of the length of the gene was composed of a specific node category, the gene was classified as that category. The models of sequence, gene and effectors were obtained by the iteration of different combinations of genomes in the sequence graph. The nodes and the genes were reclassified at each combination of genomes and the results were plotted as boxplots to include the variation between different genome combinations. To visualize the trends per category smoothed lines were applied with a smoothing span of 0.8 and 95% confidence intervals.

### Syntenic orthology inference

The primary genes of *P. theobromicola* MB01960 and P0449 plus the genes of the 9 other *Phytophthora* species (**Supplementary table 4**) with different hosts, including cacao, were used to perform the syntenic orthology analysis. The CDS and bed files of these species were processed with GENESPACE v.1.3.1 (Lovell et al. 2022). After running the main pipeline the function “syntenic_pangenes” was applied iteratively using each genome as reference. The genes with a flag PASS were count in each genome to obtain binary matrix with the refence gene and presence absence in all the other genomes. Genes with the flag PASS in at least one other genome and not with itself was counted as syntenic ortholog (SynOr). A gene was classified as SynOr conserved in all *Phytophthora* species if they were present in all the genomes (including at least one *P. theobromicola* isolate). A gene was classified as SynOr conserved only in cacao pathogens if they were present in all cacao pathogens (*P. theobromicola, P megakarya, P. palmivora, P. capsici* and *P. citrophthora*) and no more than three other species. And a gene was considered a SynOr conserved only in *P. theobromicola* in they are only present in the *P. theobromicola* genomes. The effector annotation was intersected with this information to obtain the functions of the effectors in these SynOr categories.

### RxLR homology network analysis

The total set of putative RxLR effectors in the cacao pathogens were compared against each other using the blastp of Diamond v2.1.9.163 (Buchfink, Reuter, and Drost 2021) with the options “--evalue 1e-10 --max-target-seqs 0 --outfmt 6 qseqid sseqid pident length evalue bitscore”. The result file was filtered to keep only matches with at least 30% of coverage and at least 50% identity. The network was visualized with Gephi v.0.1.0 (Bastian, Heymann, and Jacomy 2009) using the bitscore as edges. The network was arranged using the Fruchterman-Reingold layout (Fruchterman and Reingold 1991) and the cluster were defined by the modularity class calculated within Gephi. The color annotation was done using the SynOr classification. The homologous proteins of each clusters with SynOr conserved only in *P. theobromicola* were aligned with Muscle v.5.3 (Edgar 2022) and Maximum Likelihood (ML) trees were crated for each cluster using RAxML-NG v.0.9.0 (Kozlov et al. 2019) with the options “--tree pars{10} --bs-trees 100 --model WAG+I+G4”. The trees were visualized and annotated with the R packages Ape v.5.8-1 (Paradis and Schliep 2019), phytools v.2.4-4 (Revell 2024) and ggtree v.3.14.0 (Xu et al. 2022).

### Gene duplication classification

The proteins sequences of each *P. theobromicola* primary assembly were blasted against each other using Diamond v2.1.9.163 with the options “evalue 1e-3 --outfmt 6”. Matches with at least 50% coverage and identity were kept. These results passed through the MCScanX v.1.0.0 pipeline (Yupeng Wang et al. 2012) and the program “duplicate_gene_classifier” distributed with MCScanX was used to classify the observed duplication as segmental (match genes in syntenic blocks), tandem (continuous repeat), proximal (close region but not adjacent), dispersed (none of the previous categories) and singleton (no duplication). This classification was intersected with the SynOr classification and the effector annotation to extract further insights.

### Transposable element proximity and intergenic space

The gene coordinates of the primary assemblies *P. theobromicola* MB01960 and P0449 were concatenated and sorted. Similarly, the Class I and Class II TE coordinates of both genomes were concatenated and sorted (one file per class type). The tool “closest” within BEDtools v2.29.1 (Quinlan 2014) with the options “-d -io” was used to determine the distance of each gene to the closest TE on each class. Then, for each gene, the closest TE (of any kind) was selected, and the distance was recorded. Next, the genes were grouped per effector functional annotation, and a Kolmogorov–Smirnov test was used to compare the distributions of TE proximity between gene groups. A significant result indicates that one group is consistently closer or further from TEs than the other group. This information was intersected with the gene duplication classification and effector annotation to gain more insights.

The intergenic space was calculated using the tool “closest” within BEDtools v2.29.1 with the coordinates of all the genes and the options “-D a -io -k 2”. The distance to the closest gene was recorded on both sides of the genes when possible. Genes with only one closest gene were located at the end of the scaffolds. This information was intersected with RxLR effector annotation and SynOr categories to gain further insights.

### Expression analysis

RNAseq paired-end reads of all samples (mycelium, inoculated tissues and control tissues) trimmed with Trimmomatic v0.36 (Bolger, Lohse, and Usadel 2014) with the options “LEADING:7 TRAILING:7 SLIDINGWINDOW:4:15 MINLEN:50”. The genome of *Theobroma cacao* (GCF_000208745.1; (Argout et al. 2017) was concatenated with the primary assembly of *P. theobromicola* P0449 to use as reference for mapping. The reads were mapped using the Nextflow v.24.10.2 pipeline nf-core/rnaseq (v3.18.0-gb96a753; https://github.com/nf-core/rnaseq/; 10.5281/zenodo.1400710) with the options “pseudo_aligner: salmon, extra_salmon_quant_args: --seqBias --gcBias”. The transcripts and gff files were also a concatenation of both cacao and pathogen. Estimation and statistical analysis of expression level using the gene_counts_length_scaled data from salmon were performed using the DESeq2 v.1.46.0 (Love, Huber, and Anders 2014) in R. The normalized counts in inoculated tissue and control tissue were calculated within DESeq2 using the size factors account for differences in library depth (**Supplementary table 5**). The differential expressions (DE) analysis between infected tissues was calculated with the default parameters of DESeq. The contrasts of each pair of tissues were extracted and the thresholds for DE were set to Log2FC larger than 1 and adjusted p-value lower than 0.05. Upregulated genes on each tissue were extracted from the contrast with the other 2 tissues to create an upset plot. The **Supplementary table 6** present the upregulated genes and their classification per tissue type. Last, the DE genes between in-planta (inoculated tissue) and in-vitro (mycelium) were calculated using only the pathogen transcripts in the inoculated dataset. The datasets were processed with the default parameters of DESeq including the size factors normalization to account for differences in library depth. The DE genes were selected based on the same thresholds as before. The in-planta upregulated genes were binned into windows of Log2FC of five units (from 1 to 5, from 5 to 10, from 10 to 15 and so on) for figure preparation. The full table of DE genes between in-planta vs in-vitro is presented in **Supplementary Table 7**.

## Results

### Genome assembly of Phytophthora theobromicola

The genomes of two *P. theobromicola* isolates, MB01960 and P0449, collected in Bahia, Brazil, were assembled using PacBio CLR long-read sequencing (**Table 1**). Each assembly included both primary contigs and associated haplotigs, which were uniformly distributed across the genome, consistent with a diploid representation. The haploid genome sizes were estimated at 47.5 Mbp for MB01960 and 45.6 Mbp for P0449, with sequencing coverage exceeding 700 X. These genome sizes place *P. theobromicola* among the smaller cacao-infecting *Phytophthora* species, similar to *P. citrophthora*, and considerably smaller than *P. capsici* and *P. palmivora* (approximately half), and *P. megakarya* (less than one-quarter). Genome completeness, assessed using BUSCO on both genomic sequences and predicted proteomes, was high for both isolates, exceeding 96%. Predicted gene counts followed a pattern consistent with genome size, showing similar values to those observed in *P. citrophthora*. In addition, five more *P. theobromicola* isolates were sequenced using short-read technologies and *de novo* assembled (**Supplementary Table 1**). These assemblies had an average size of 43.8 ± 0.3 Mbp (considering scaffolds >1 kb), with coverage above 400 X. The average genome completeness for these assemblies was 88.8 ± 2.0%, and the number of predicted genes was approximately 32% lower than in the long-read assemblies, likely reflecting limitations in assembly contiguity and completeness using short-read sequencing.

### Effector identification and functional annotation

*Phytophthora* species, like other oomycete pathogens, are known to secrete a large repertoire of effector proteins that facilitate host colonization and suppress plant defenses (McGowan and Fitzpatrick 2017; Wawra et al. 2012). We functionally annotated the putative effector repertoires of the five cacao-infecting *Phytophthora* species listed in **Table 1**, including both *P. theobromicola* isolates. To identify secreted proteins, we first predicted the secretome of each species using SignalP3 in combination with transmembrane domain prediction, retaining only proteins with a secretion signal and no transmembrane domains outside the signal peptide. Proteins with predicted secretion signals and domain annotations consistent with known effector categories were classified as putative secreted effectors. From a total of 157,650 predicted proteins across the primary assemblies of all species, 12,631 were identified as putatively secreted. On average, 44.2 ± 2.0% of the secretome per species was annotated as putative effectors (**Table 2**). In *P. theobromicola*, RxLR effectors and CAZymes together accounted for more than 67% of the predicted effectors, while in *P. megakarya*, these two categories comprised over 78%, with RxLRs alone representing 56.7% (**Table 2**). Among the CAZymes, we identified numerous enzymes implicated in cell wall degradation, including those targeting pectin, lignin, and hemicellulose. Additionally, several elicitins, NLPs, and phytotoxins were detected in the secretome of all species. A full list of annotated effector genes, including those from the short-read assemblies, is provided in **Supplementary Tables 2&3**. Summary statistics by effector category are available in **Supplementary Table 8**.

**Table 2.** Counts of effectors proteins classified per category or function in the species in study.

| Category | *P. theobromicola*<br>(MB01960) | *P. theobromicola*<br>(P0449/ATCC) | *P. megakarya*<br>(PmGH34) | *P. palmivora*<br>(PpGH49) | *P. capsici*<br>(LT1534) | *P. citrophthora*<br>(STE-U-9442) |
| --- | --- | --- | --- | --- | --- | --- |
| Number of secreted proteins | 1,718 | 1,628 | 3,541 | 2,623 | 1,939 | 1,182 |
| Number of effectors | 695 | 638 | 1,807 | 1,252 | 823 | 522 |
| Effector proportion (%) | 40.5 | 39.2 | 51.0 | 47.7 | 42.4 | 44.2 |
| RxLR | 219 | 208 | 1,026 | 593 | 295 | 214 |
| Crinkler | 3 | 2 | 0 | 0 | 3 | 3 |
| CAZymes | 253 | 223 | 395 | 370 | 313 | 167 |
| Glycoside hydrolases | 127 | 126 | 202 | 212 | 150 | 86 |
| Cellulases | 28 | 25 | 44 | 45 | 32 | 16 |
| Chitinases | 3 | 3 | 2 | 5 | 4 | 6 |
| Cutinases | 6 | 6 | 11 | 7 | 9 | 4 |
| Hemicellulose degradation | 28 | 27 | 57 | 62 | 38 | 24 |
| Lignin degradation | 10 | 9 | 13 | 6 | 8 | 4 |
| LPMOs | 65 | 47 | 87 | 92 | 73 | 25 |
| Pectin degradation | 66 | 63 | 113 | 94 | 81 | 59 |
| Elicitins | 51 | 44 | 47 | 54 | 39 | 42 |
| NLP | 57 | 47 | 147 | 113 | 43 | 28 |
| Phytotoxins | 5 | 5 | 0 | 1 | 5 | 6 |
| Trypsin & Trypsin like proteins | 22 | 23 | 56 | 24 | 25 | 13 |
| Kazal-type Protease inhibitors | 18 | 17 | 72 | 37 | 23 | 17 |
| Papain and papain like family cysteine proteases | 12 | 12 | 10 | 10 | 21 | 6 |

### Sequence graph analysis reveals high intraspecific variability of the effector repertoire in *P. theobromicola*

Genomic variability among isolates of a single species has been shown to significantly impact the composition and diversity of virulence factors in plant pathogens, contributing to their ability to adapt to diverse environmental conditions (Garcia et al 2023). Analyses of *P. palmivora* and *P. megakarya* revealed that effector genes particularly those belonging to the RxLR family are overrepresented among genes under positive selection (Morales-Cruz et al., 2020). To assess intraspecific effector variability in *P. theobromicola*, we constructed a sequence graph that integrates the genomic content of all available isolates. This approach enables the identification of both conserved and variable regions across genomes and allows for classification of effector genes into three categories: core (present in all isolates), dispensable (shared by more than one but not all), and private (unique to a single isolate). The graph was constructed using primary contigs and haplotigs from the two long-read assemblies, along with the five short-read genome assemblies, resulting in a combined graph spanning 67.5 Mbp (**Figure 1A**). This represents an 83% compression relative to the ∼400 Mbp cumulative length of the individual assemblies, reflecting a high degree of shared sequence content and capturing structural variation across isolates.

**Figure 1.**
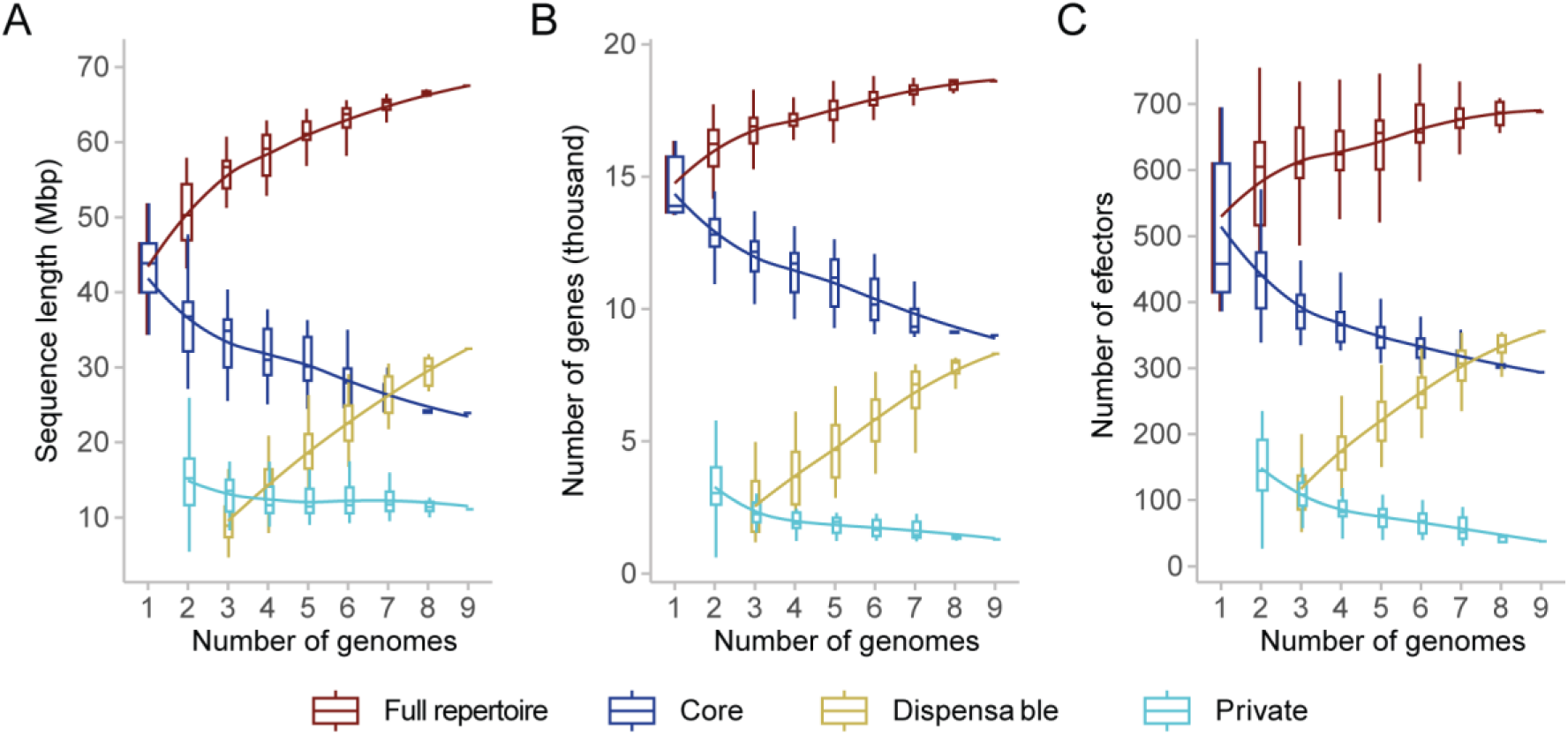
Variability within the *P. theobromicola* species represented as sequence graph models of A) sequence length, B) Number of total genes and C) number of effectors. All panels share the same color legend.

After classifying the genes of each isolate based on their position in the graph (core, dispensable, or private), we observed a linear increase in the number of dispensable genes with each additional genome included (**Figure 1B**). This trend persisted until the inclusion of the eighth genome, after which the slope began to decrease slightly, suggesting that the set of dispensable genes may be approaching saturation. A similar pattern was observed when focusing specifically on effector genes: the number of dispensable effectors increased linearly and steeply across genomes, while the total effector repertoire appeared to begin plateauing after the third genome was added. A detailed analysis of the final effector graph revealed that, on average, 352 ± 30 effector genes per genome were classified as core, representing 68.0 ± 1.5% of the total effector repertoire. Dispensable effectors accounted for 31.6 ± 1.5%, and only a small fraction, 0.4 ± 0.1%, were classified as private (**Supplementary Figure 1A&B**). These results indicate substantial intraspecific variability in effector content, while also suggesting the presence of a relatively stable core set across isolates.

### Comparative syntenic orthology reveals lineage-specific effector conservation in *P. theobromicola* and other cacao pathogens

To investigate patterns of gene conservation and diversification within *P. theobromicola*, we performed a syntenic orthology analysis using GENESPACE across 13 *Phytophthora* species, including the two long-read *P. theobromicola* primary assemblies (**Supplementary Table 4**). By incorporating the genomic context of orthologous groups, this approach enabled the detection of conserved gene blocks while accounting for structural rearrangements across genomes. Genes located within syntenic regions were classified into three categories: those conserved across all species, those conserved only among cacao-infecting species, and those unique to *P. theobromicola*.

Over 80% of genes in *P. theobromicola* have syntenic orthologs in at least one other *Phytophthora* species (**Figure 2A**). Additionally, approximately 9% of the total gene content corresponds to local duplications of syntenic orthologs (**Figure 2B**). These proportions contrast with species such as *P. megakarya* and *P. fragariae*, which have genome sizes and gene counts more than 2.5 times greater than those of *P. theobromicola*. From the total syntenic gene sets analyzed, an average of 40.8 ± 0.2% of genes are conserved across all species (**Figure 2C**). In contrast, only 1.4 ± 0.02% are conserved exclusively among cacao-infecting *Phytophthora* species (**Figure 2D**), while 5.8 ± 0.02% appear to be conserved only in *P. theobromicola* (**Figure 2E**). These results highlight the presence of both highly conserved genes and a subset of lineage-specific genes that may contribute to host adaptation and species-specific functions.

**Figure 2.**
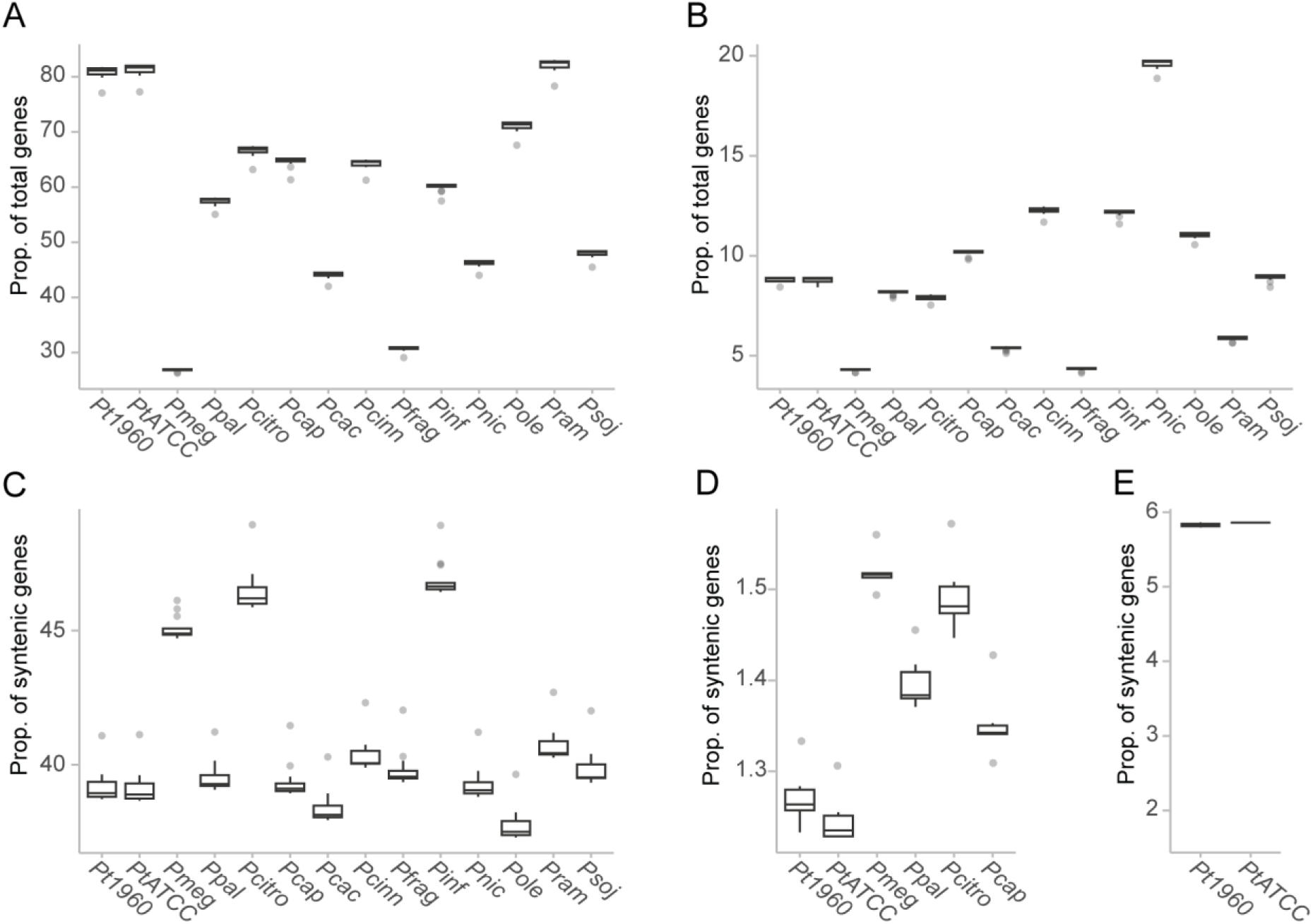
Syntenic orthologous gene groups in multiple *Phytophthora* species. A) Proportion of total genes with syntenic orthologs in at least one other species. B) Proportion of total genes derived from duplication of a syntenic ortholog. C) Proportion of syntenic orthologs conserved in all *Phytophthora* species in study. D) Proportion of syntenic orthologs conserved only in cacao pathogens. E) Proportion of syntenic orthologs conserved only in *P. theobromicola*. Pt1960: *P. theobromicola* MB01960, PtATCC: *P. theobromicola* P0449/ATCC, Pmeg: *P. megakarya*, Ppal: *P. palmivora*, Pcitro: *P. citrophthora*, Pcap: *P. capsici*, Pcac: *P. cactorum*, Pcinn: *P. cinnamomic*, Pfrag: *P. fragariae*, Pinf: *P. infestans*, Pnic: *P. nicotianae*, Pole: *P. oleae*, Pram: *P. ramorum* and Psoj: *P. sojae*.

To further examine gene variability across *P. theobromicola* isolates, we intersected the syntenic ortholog categories with the graph-based classifications and effector annotations. This integration enabled us to assess how conserved and lineage-specific genes contribute to the core and dispensable genomes, with a particular focus on effectors. On average, 3,809 ± 31 syntenic orthologs (SynOrs) conserved across all *Phytophthora* species were also classified as core genes in *P. theobromicola*, representing 35.3% of the species’ core genome based on the sequence graph (**Supplementary Figure 2A**). In addition, 1,616 ± 61 universally conserved SynOrs were found in the dispensable category, comprising ∼31% of the dispensable gene set per genome. Among SynOrs conserved exclusively in cacao-infecting *Phytophthora* species, 133 ± 1 were core genes (1.2% of the total core genome), and 77 ± 9 were dispensable (1.5% of the dispensable set; **Supplementary Figure 2B**). Similarly, SynOrs conserved only in *P. theobromicola* accounted for 4.4% of the total core genes and approximately 6.7% of the dispensable genes (Supplementary Figure 2D). Effector genes followed similar patterns. On average, 9.0 ± 0.2% of effectors per genome were classified as SynOrs conserved across all species (**Supplementary Figure 3A**), 2.5 ± 0.1% were conserved only among cacao pathogens (**Supplementary Figure 3B**), and 4.1 ± 0.03% were conserved exclusively in *P. theobromicola* (**Supplementary Figure 3C**). Within these groups, RxLR effectors and CAZymes were particularly well represented (**Figure 3A&B**), alongside other functional categories such as cutinases, hemicellulose-degrading enzymes, LPMOs, NLPs, and phytotoxins.

**Figure 3.**
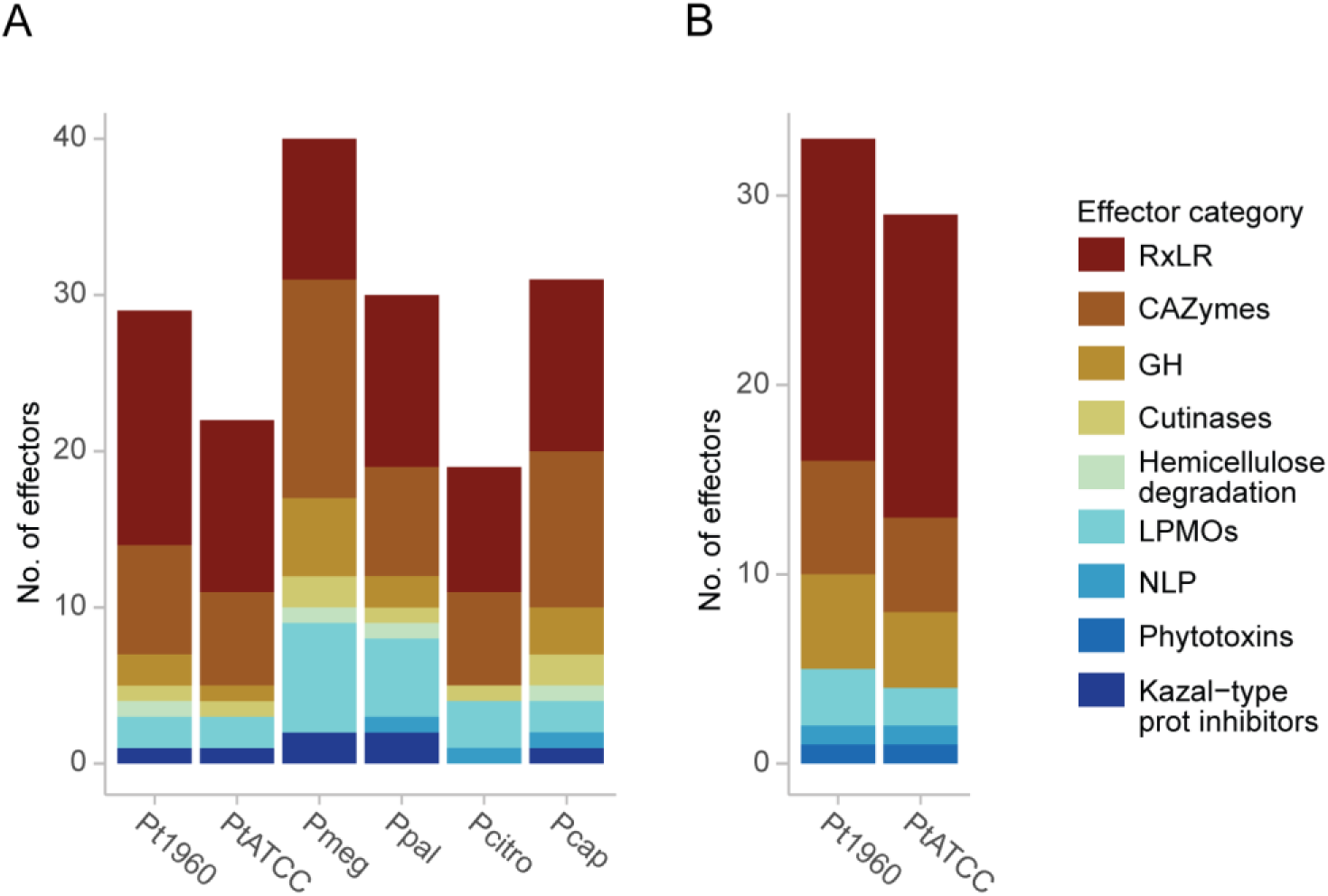
Number and function of effectors classified as syntenic orthologs A) conserved only in cacao pathogens and B) conserved only in *P. theobromicola*. Panels A and B share the same color legend. Pt1960: *P. theobromicola* MB01960, PtATCC: *P. theobromicola* P0449/ATCC, Pmeg: *P. megakarya*, Ppal: *P. palmivora*, Pcitro: *P. citrophthora*, Pcap: *P. capsici*.

### Analysis of homology, duplication, and intergenic space highlights genomic drivers of effector variation

To investigate the evolutionary relationships and diversification patterns of RxLR effectors among cacao-infecting *Phytophthora* species, we performed a homology-based network analysis. All predicted RxLR protein sequences were compared using BLASTp, and the resulting similarity network was visualized and clustered using Gephi. This approach allowed the grouping of RxLRs into putative families based on shared sequence features, while also highlighting their SynOr categories as defined in the previous analyses. Several *P. theobromicola*-specific SynOrs clustered with RxLRs from multiple species, as well as with SynOrs conserved across broader taxonomic groups (**Figure 4**). For each of the highlighted clusters in the network, protein sequences were aligned using MUSCLE (v5.3), and maximum likelihood phylogenetic trees were constructed (**Supplementary File 1**). Within each tree, SynOrs conserved exclusively in *P. theobromicola* consistently grouped together, forming distinct clades. The closest homologs to these SynOrs typically originated from *P. citrophthora* or *P. capsici*, and rarely from *P. megakarya* or *P. palmivora*. This pattern suggests that the observed variability may arise from lineage-specific duplications, gene rearrangements, or divergence from a shared ancestral gene set.

**Figure 4.**
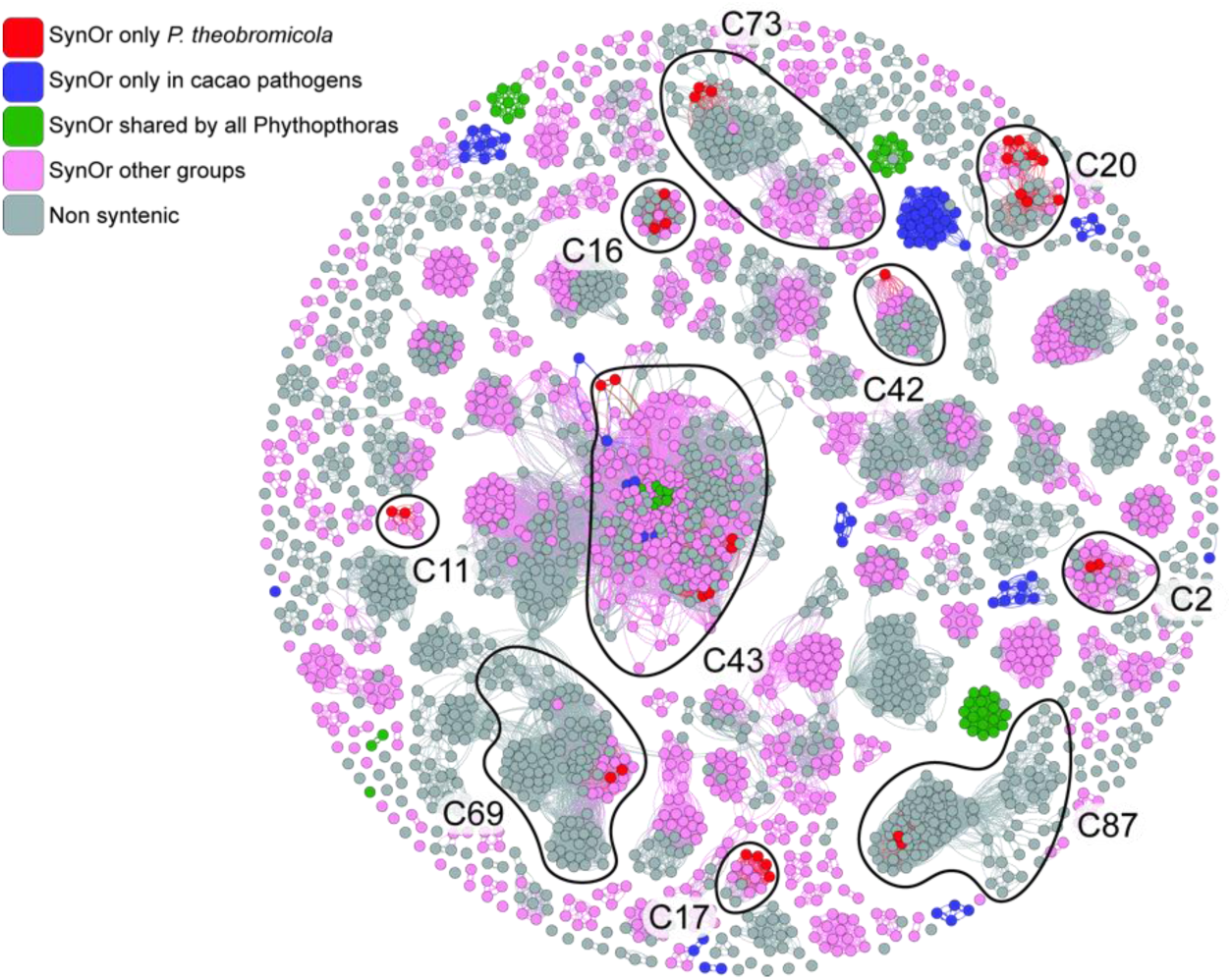
Homology network analysis of all the putative RxLR effectors among cacao-infecting *Phytophthora* species. Each RxLR is represented by a node. Edges connection between two nodes represent similarity shared by the by the two proteins. Clusters were calculated the interconnectedness between proteins as described in methods. Nodes are colored based on the syntenic orthology groups.

Given the role of transposable elements (TEs) in promoting genome plasticity and effector diversification (Raffaele et al. 2010; Morales-Cruz et al. 2020), we examined the spatial distribution of RxLRs and other effector categories in relation to annotated Class I and Class II TEs in *P. theobromicola*. Genes located near TEs are frequently subject to duplication, insertional mutagenesis, and other forms of structural variation (Faino et al. 2016; Schrader and Schmitz 2018; Dong, Raffaele, and Kamoun 2015). In our analysis, RxLR effectors were found to be significantly closer to TEs than any other effector category, with the exception of NLPs (Kolmogorov–Smirnov test, *p* < 0.016; **Figure 5A**).

**Figure 5.**
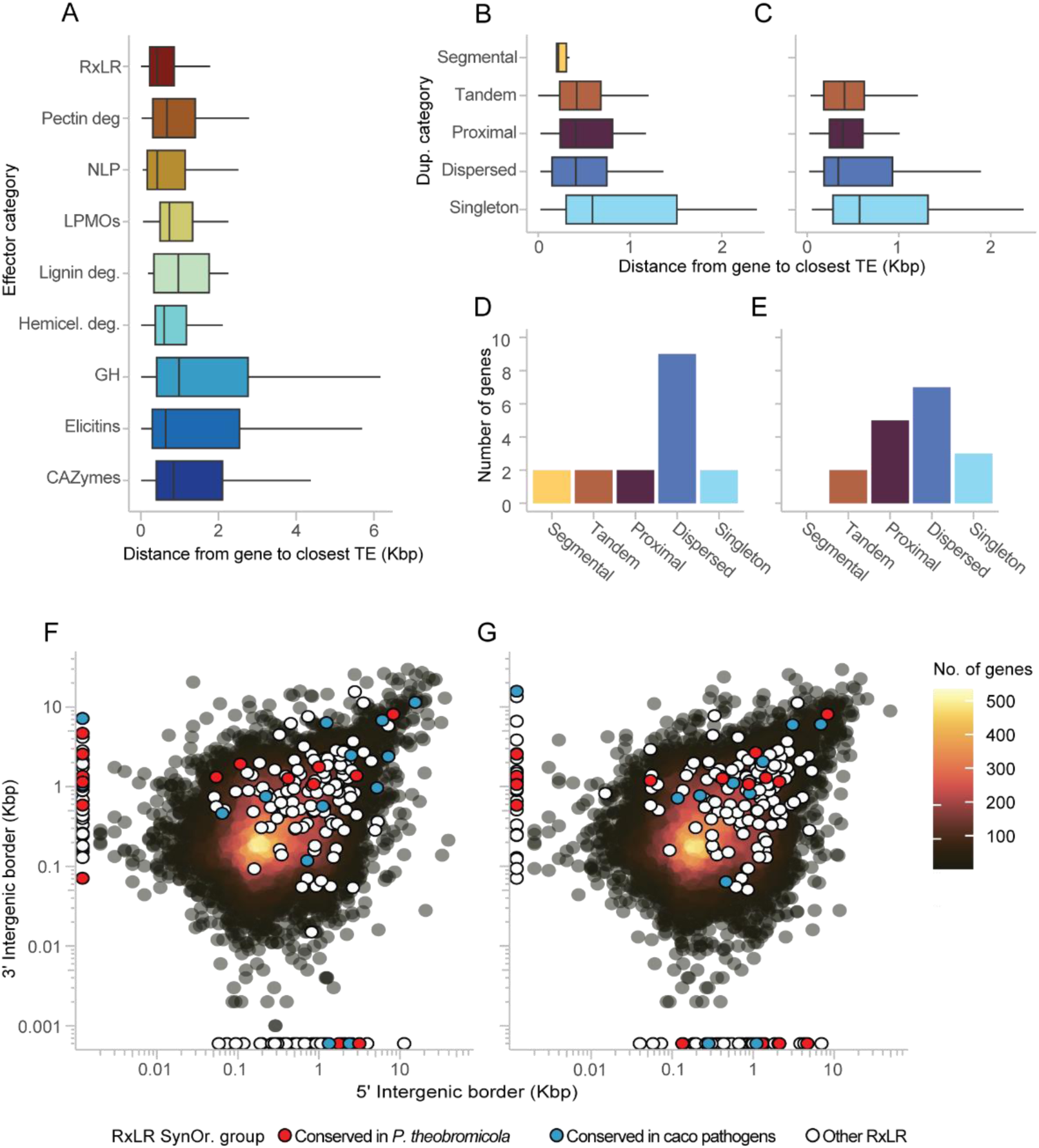
Classification of *P. theobromicola* effectors based on proximity to transposable elements (TEs) and intergenic space. A) Distance of different effector functional categories to the closest annotated TE. B–C) RxLR effectors classified by duplication category and their distance to the nearest TE in isolates MB01960 and P0449, respectively. D–E) Number of SynOr RxLRs conserved only in *P. theobromicola*, classified by duplication category in isolates MB01960 and P0449, respectively. F–G) Intergenic space surrounding genes in isolates MB01960 and P0449, respectively, highlighting RxLR effectors and colored by SynOr group classification. Panels F and G shares the same color legend.

To investigate the origins of RxLR variability, we conducted a duplication classification using MCScanX and assessed the proximity of each RxLR gene to nearby TEs. Across isolates, *P. theobromicola* harbored an average of 214 ± 6 RxLR effectors, of which 6 were derived from segmental duplications, 47 ± 2 from tandem duplications, 52 ± 2 from proximal duplications, 42 ± 1 from dispersed duplications, and 71 ± 2 were singletons. There were no statistically significant differences in the distribution of duplication categories between isolates (Kolmogorov–Smirnov test); however, segmental duplications were only detected in the MB01960 isolate (**Figure 5B&C**). Notably, most of the *P. theobromicola*-specific SynOr RxLRs were associated with dispersed and proximal duplications, whereas those classified as singletons frequently mapped near DNA transposable elements (**Figure 5D&E**). These findings support a role for both local duplication and TE-mediated mechanisms in shaping RxLR diversity in this species.

To further explore the sources of variability in the effector repertoire of *P. theobromicola*, we quantified intergenic distances surrounding all genes, with a focus on RxLR effectors. Larger intergenic regions have been associated with relaxed selective constraints and increased potential for sequence diversification through mutation or structural rearrangement rearrangements (Raffaele and Kamoun 2012; Dong, Raffaele, and Kamoun 2015). RxLR genes exhibited significantly larger intergenic distances compared to all other genes (Wilcoxon rank-sum test, *p* < 2.2 x 10⁻¹⁶), with a mean intergenic distance of 1,391 ± 71 bp, in contrast to 697 ± 6 bp for non-RxLR genes (**Figure 5F&G**). Furthermore, SynOr RxLRs conserved exclusively in cacao-infecting species and those unique to *P. theobromicola* displayed similar intergenic lengths, both significantly greater than those of the remaining RxLRs (Dunn’s test, *p* < 0.01; **Figure 5F&G**). These results suggest that the broader intergenic context of these effectors may facilitate their diversification, particularly for lineage-specific RxLRs, reinforcing the role of genomic architecture in shaping effector evolution.

### Expression profiling reveals in-planta activation of effectors including the lineage-specific

While gene prediction provides a useful overview of the genomic potential of a species, expression profiling helps determine which genes are potentially active under biologically relevant conditions. To assess the *in planta* expression of *P. theobromicola* genes during infection, we inoculated three different cacao tissues, pods, leaves, and stems, with mycelial plugs from the P0449 isolate and performed RNA-seq analysis alongside matched uninoculated controls. Among the infected tissues, pods exhibited the highest number of expressed *P. theobromicola* genes, followed by leaves and stems (**Supplementary Figure 4A-C**). About 88% of the predicted effector repertoire was expressed in at least one tissue. Pods showed the highest expression across all effector categories, with approximately 70% of predicted RxLRs, 100% of Crinklers, and 94% of effector-associated CAZymes detected in the transcriptome (Supplementary Figure 5).

Differential expression analysis revealed tissue-specific responses. A total of 9,300 *P. theobromicola* genes were significantly upregulated in pods relative to both leaves and stems (log₂FC > 1; adjusted *p*-value < 0.05). Additionally, 1,477 genes were upregulated in both pods and leaves compared to stems, and 1,041 genes were specifically upregulated in leaves. In contrast, very few genes were preferentially expressed in stems (**Figure 6**). Among the genes upregulated exclusively in pods, 89 were RxLR effectors, including three classified as SynOrs conserved only in *P. theobromicola* and six conserved only among cacao-infecting species (**Figure 6**). These results suggest tissue-specific effector deployment, with pods being a major site of effector activity during infection.

**Figure 6.**
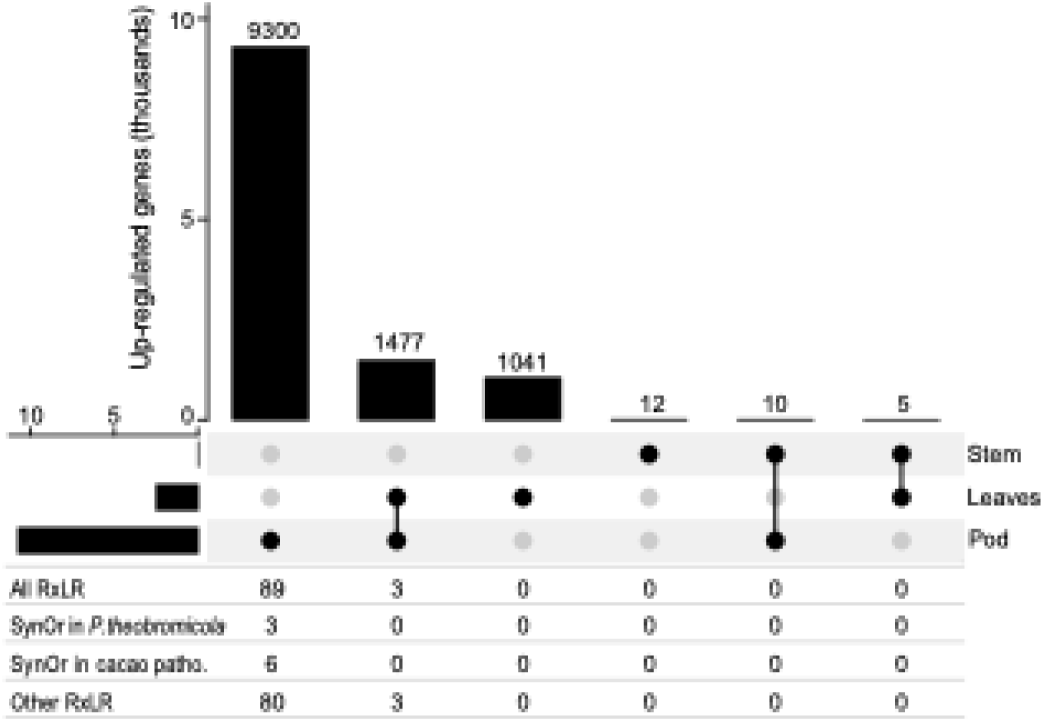
Comparison of up-regulated genes per inoculated tissue. The table present the number RxLR effectors in each bar including the total set of RxLR and the RxLRs in the different SynOr categories.

Gene expression in inoculated tissues was compared to pure mycelial cultures to identify differentially expressed (DE) genes. A total of 5,142 genes were differentially expressed, with 63% upregulated *in planta* (**Figure 7A**), including 46% of the predicted effector repertoire (**Figure 7B**). Among these, RxLR effectors and CAZymes showed a high proportion of genes with log₂ fold changes (L2FC) between 1 and 5 (**Figure 7C**). Interestingly, the number of RxLR genes with L2FC values between 5 and 10 was nearly double that of the previous range and included two SynOrs conserved exclusively in *P. theobromicola* (**Figure 7D**). Two out of the three *P. theobromicola*-specific SynOr RxLRs classified as singletons were strongly upregulated *in planta*, showing 5-fold and 53-fold higher expression, respectively, compared to *in vitro* mycelial samples. A smaller subset of effector genes exhibited even more extreme differential expression, with L2FC values between 10 and 15 (**Figure 7E**). Among these, one of the five phytotoxins encoded in the genome showed over 1,000-fold higher expression *in planta*. Additionally, a SynOr RxLR conserved only among cacao-infecting species was expressed 4,500 times more *in planta* than *in vitro*. These findings suggest strong host-induced expression of key effectors, including lineage-specific genes that may play critical roles in pathogenicity and host adaptation.

**Figure 7.**
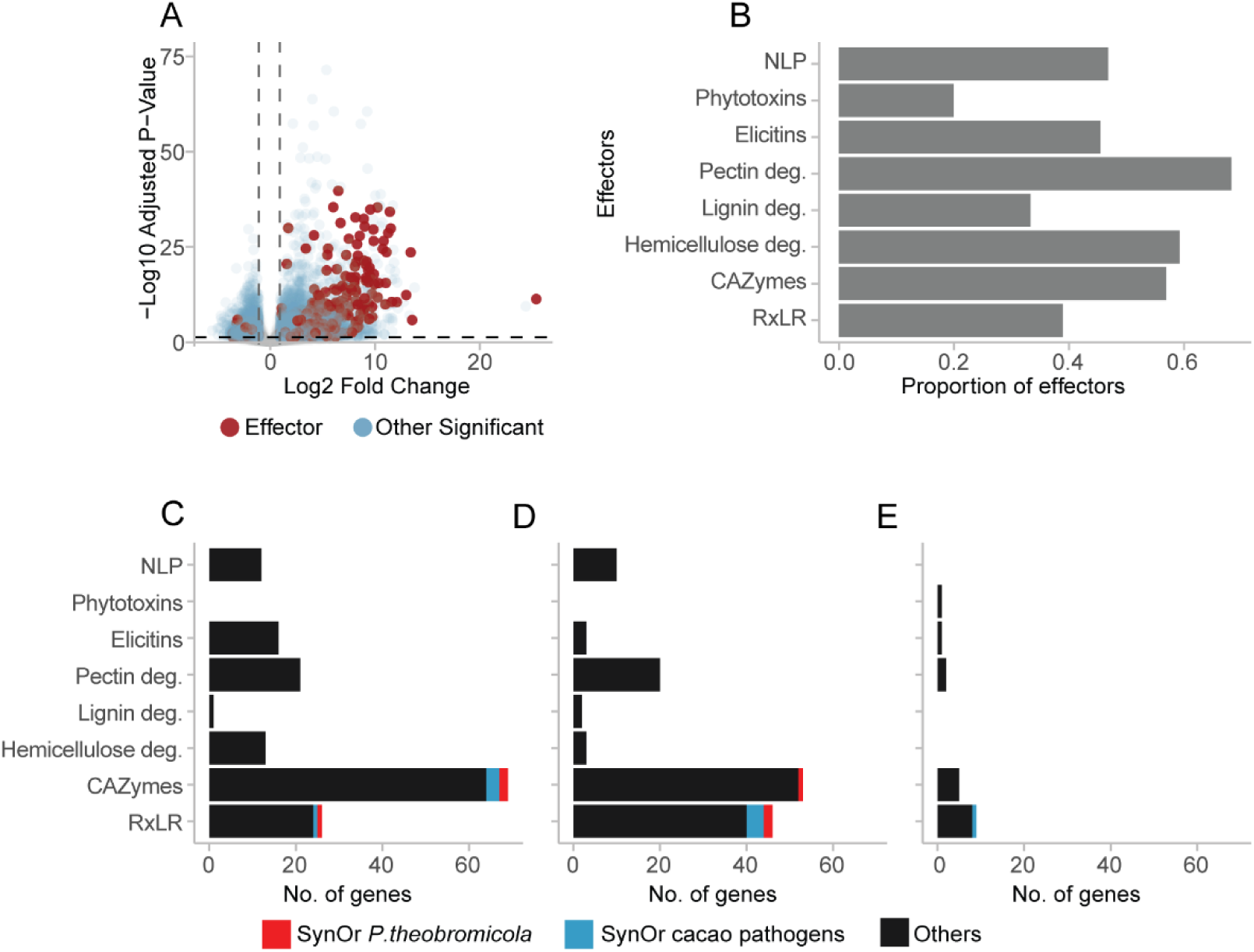
Differential expression of P*. theobromicola* genes *in planta* compared to *in vitro*. A) Volcano plot showing differentially expressed genes between *in planta* and *in vitro* conditions, highlighting effectors. B) Proportion of total effectors from each functional category upregulated *in planta*. C–E) Number of genes with Log2 fold change (L2FC) values between 1-5 (C), 5-10 (D), and 10-15 (E). Panels C–E share the same color legend.

## Discussion

*Phytophthora theobromicola* has recently emerged as a major contributor to black pod rot disease in Brazil, exhibiting levels of aggressiveness comparable to those of *P. palmivora* (Decloquement et al. 2021). Here, we report high-quality and high-completeness genome assemblies generated from multiple *P. theobromicola* isolates. The assembled genome size and number of predicted protein-coding genes are similar to those of its close relative *P. citrophthora*. This finding is consistent with earlier reports of misidentification of *P. theobromicola* as *P. citrophthora* because of their morphological similarities (Luz et al. 2018; Decloquement et al. 2021).

Access to high-quality genome assemblies enabled the annotation of putative effector genes in *P. theobromicola*. Approximately 40% of the predicted secretome in the assembled genomes was assigned to known effector categories. While this proportion is comparable to that observed in other *Phytophthora* species infecting cacao, the total number of effectors in *P. theobromicola* is considerably lower than in species such as *P. megakarya* and *P. palmivora*. This discrepancy is partially explained by the whole-genome duplications previously reported in *P. megakarya* and *P. palmivora*, which resulted in expanded gene content and two of the largest effector repertoires within the genus (Morales-Cruz et al. 2020).

Nonetheless, the repertoire of virulence factors, and particularly effectors, can vary substantially among isolates, as demonstrated in previous studies of filamentous pathogens (Morales-Cruz et al. 2020; Garcia et al. 2024). To better capture this intraspecific variation, we constructed a sequence graph that integrates the genomes of multiple *P. theobromicola* isolates, enabling a more comprehensive view of the species’ effector complement. The marked increase in the proportion of dispensable genes and effectors, along with the finding that over 30% of predicted effectors are dispensable, indicates a high degree of genomic variability in *P. theobromicola*. Similar patterns have been reported in other highly virulent filamentous pathogens (Garcia et al. 2024). The relatively slow growth of the total effector repertoire with the addition of new genomes (**Figure 1 C**) suggests that while novel genomes continue to contribute new genes, their greatest impact is on the dispensable and core proportions, a pattern likely driven by the intrinsic genomic plasticity of *Phytophthora* species (Kronmiller et al. 2023).

Syntenic orthology analysis provided a framework to further explore genomic flexibility and track patterns of effector diversification in *P. theobromicola*. By comparing a diverse set of *Phytophthora* species with varying host ranges, we identified syntenic ortholog groups conserved across all species, conserved only among cacao-infecting pathogens, and unique to *P. theobromicola*. Within the effector genes falling into cacao-specific and *P. theobromicola*-specific SynOr categories, RxLRs and CAZymes were particularly enriched. RxLR effectors, in particular, have been consistently associated with expanded intergenic regions and elevated evolutionary rates compared to other genomic regions in *Phytophthora* (Morales-Cruz et al. 2020; Dong, Raffaele, and Kamoun 2015; Raffaele et al. 2010; Stassen and Van den Ackerveken 2011). These characteristics likely contribute to the high variability in orthology and disrupted collinearity observed even between species within the same genus.

The relationship between RxLR effectors and SynOr gene classification was further explored through a homology-based network including RxLR sequences from multiple cacao-infecting *Phytophthora* species. Clustering patterns and phylogenetic relationships within these groups suggest that SynOrs conserved exclusively in *P. theobromicola* likely originated through lineage-specific duplication or translocation events. Previous studies have shown that such genomic variability is often associated with the activity of transposable elements (TEs), which can drive effector diversification and structural genome variation (Raffaele and Kamoun 2012; Faino et al. 2016; Schrader and Schmitz 2018; Morales-Cruz et al. 2020; Zhang et al. 2024). Consistent with this, RxLRs and a subset of other effectors in *P. theobromicola* are located significantly closer to TEs than CAZymes or non-effector genes, mirroring patterns previously reported in other *Phytophthora* pathogens of cacao (Morales-Cruz et al. 2020).

Moreover, SynOr RxLRs conserved only in *P. theobromicola* that do not arise from duplication events (i.e., singletons) are frequently located near DNA transposable elements. Given that DNA elements typically follow a cut-and-paste mechanism (Wicker et al. 2007), these RxLRs were likely translocated from other genomic regions (Schrader and Schmitz 2018; Raffaele and Kamoun 2012), resulting in structural rearrangements unique to *P. theobromicola*. The larger intergenic distances observed around RxLR genes, compared to other genes, are consistent with the two-speed genome model, in which effector genes, particularly RxLRs, reside in gene-sparse, repeat-rich regions that evolve more rapidly and contribute to host adaptation (Dong, Raffaele, and Kamoun 2015; Frantzeskakis, Kusch, and Panstruga 2019). Notably, many RxLRs were located near scaffold ends, a pattern likely caused by large intergenic, repeat-rich regions associated with RxLRs that complicate assembly and frequently result in contig breaks.

Gene expression analysis of *P. theobromicola* in different inoculated cacao tissues revealed that at least 88% of the predicted effector genes are transcriptionally active, including 70% of the total predicted RxLRs expressed in pods. This proportion of active RxLRs is comparable to that observed in *P. megakarya* and *P. palmivora*, where approximately 75% of RxLRs are expressed *in planta* (Morales-Cruz et al. 2020). Pods appeared to be the major site of pathogen activity, as most upregulated genes were specific to this tissue. Among them were 89 RxLRs, including nine classified as SynOrs conserved exclusively in *P. theobromicola* or among cacao-infecting *Phytophthora* species. These findings highlight pods as a key infection site and suggest a specialized deployment of effectors, particularly lineage-specific RxLRs, during host colonization.

Comparison of *P. theobromicola* gene expression *in planta* and *in vitro* revealed that the majority of differentially expressed genes were upregulated during host colonization, including most predicted effectors. Only a small subset of effectors was upregulated *in vitro*, suggesting that effector expression is primarily induced by host-derived cues and likely regulated in response to plant signals (Inoue et al. 2023; Giraldo and Valent 2013). This expression pattern is consistent with the established role of effectors in modulating plant immune responses, reinforcing their functional importance during infection. Importantly, several SynOr RxLRs conserved exclusively in *P. theobromicola* were also transcriptionally active *in planta*, indicating that these lineage-specific genes are not only structurally unique but also likely contribute to pathogenicity.

This study presents the first comprehensive genomic analysis of the black pod rot pathogen *Phytophthora theobromicola*. Our results reveal a highly flexible genome, shaped by structural rearrangements and enriched for variable effector genes, many of which are closely associated with transposable elements. Integration of sequence graph analyses across multiple isolates, synteny comparisons, and transcriptomic profiling demonstrates that numerous lineage-specific RxLR effectors are actively expressed in cacao pods and are frequently located in gene-sparse, repeat-rich regions—consistent with the two-speed genome model. Together, these findings highlight the role of genome plasticity in effector diversification and potentially host adaptation, and provide a foundation for future investigations into the evolutionary dynamics and pathogenic strategies of *P. theobromicola*.

## Data Availability

The sequencing data and genome assemblies for this project are available at NCBI (https://www.ncbi.nlm.nih.gov/bioproject/PRJNA1257449) The genome assemblies and gene models produced in this study are also publicly available at Zenodo (10.5281/zenodo.15346916). Dedicated genome browser and BLAST tools are available at cacaopathogenomics.com.

## Acknowledgments

We thank Dr. Andrea Minio for his technical support and the UC Davis DNA Technologies Core for sequencing assistance.

